# Non-genetic inheritance of stochastically induced behavioral individuality in a naturally clonal fish

**DOI:** 10.64898/2026.03.31.715612

**Authors:** Ulrike Scherer, Sean M. Ehlman, David Bierbach, Ido Pen, Jens Krause, Max Wolf

## Abstract

The study of among-individual phenotypic variation arising in the apparent absence of genetic and environmental differences has recently emerged as a rapidly growing research area. Despite growing recognition of its existence and fitness relevance, it remains unknown whether signatures of such presumably stochastically induced variation can be transmitted across generations. To address this knowledge gap, we performed a two-generation behavioral screening with a naturally clonal fish: 34 genetically identical mothers and their 232 offspring were separated after birth into near-identical environments, with early-life behavior being tracked continuously at high resolution, constituting a total of ∼19,000 observation hours. We find consistent among-individual differences in behavior (i.e., activity and feeding patterns) in both mothers and offspring. Mother behavior correlates with offspring activity (but not offspring feeding): mothers that spend more time feeding produce more active offspring. We find no evidence of body size (maternal or offspring) mediating mother-offspring behavioral associations. Our study provides first evidence for the non-genetic transmission of among-individual phenotypic differences that arise in the apparent absence of genetic or environmental differences, highlighting the potential importance of this variation for evolutionary processes and the adaptive potential of populations.

## INTRODUCTION

In recent years, the study of among-individual phenotypic variation that arises even under highly standardized genetic and environmental conditions has attracted burgeoning interest in biological research (1–4). Evidence for such seemingly ‘stochastic’ variation now spans levels of biological organization from single cells to vertebrate animals and encompasses a broad array of traits, including stochasticity in mRNA and protein production, neural development and physiological as well behavioral traits (1–3,5–7). In behavioral biology, work using three powerful study systems (naturally clonal fish, isogenic fruit flies, and inbred mice) has revealed that substantial behavioral individuality (i.e., consistent among-individual differences in behaviors) emerges in traits such as activity and exploration, feeding pattern, phototaxis, and locomotor handedness – despite the absence of apparent genetic or environmental differences among individuals (8–15). Studies on naturally clonal fish further demonstrate that these seemingly stochastic behavioral differences are not merely inconsequential noise but are linked (directly or indirectly) to key fitness components: growth, reproduction, and lifespan (14,16). Together, this body of work provides compelling evidence that apparent genetic and environmental differences alone are insufficient to account for the phenotypic variation observed in biological systems, and that stochastic processes may play a far more important role in generating biologically meaningful variation than traditionally assumed. We note that, in line with the literature (1,4,7), we here use the term ‘stochastic’ in an operational sense, encompassing variation that is truly stochastic, variation that initiates stochastically but may unfold deterministically, and variation that derives from deterministic processes but cannot be predicted from readily measurable variables – and is thus treated as unexplained residual variation when controlling for genetic and environmental sources of variation as best as is currently empirically possible. Despite growing recognition of both the existence and fitness relevance of phenotypic variation arising in the apparent absence of genetic and environmental differences, it is completely unknown whether and to what extent such variation is transmitted across generations. Resolving this question is crucial, because, if stochastically induced phenotypic differences are transmitted across generations, they would constitute a previously unrecognized factor contributing to evolution and the adaptive potential of populations.

Given that stochastic phenotypic variation arises, by definition, through unpredictable processes, the possibility that such variation could be inherited may appear counterintuitive. However, while the processes that generate stochastic phenotypic differences may not themselves be predictable, the phenotypes they produce shape an organism’s internal state. Behavior, in particular, feeds back on physiology: differences in activity or feeding, for example, can alter energy balance, body condition, hormonal profiles, or epigenetic states (17–19), all of which are plausible carriers of information to the next generation. Indeed, such non-genetic inheritance is increasingly recognized as an important mechanism by which environmentally induced differences are transmitted across generations (20–22), yet nothing is known about whether and how phenotypic differences that arise under highly standardized conditions can be passed on through these mechanisms.

Here, we employ a powerful vertebrate study species - the naturally clonal Amazon molly, *Poecilia formosa* – to test the intergenerational transmission behavioral differences arising under near-identical genetic and environmental conditions. We performed a large-scale, two-generation behavioral screening - consisting of a total of N = 34 genetically identical mothers and their N = 232 offspring - reared in near-identical environments. Mothers were separated directly after birth into standardized environments where each mother was recorded for 10 hours per day over the first 28 days of life. The emerging among-individual differences in activity profiles and feeding behavior were characterized at high resolution (5 datapoints per second for 10 hours per day over 28 days, amounting to roughly 171 million datapoints and 9,520 recording hours) (23). Once behavioral assays were completed, individuals were allowed to reproduce in standardized, near-identical housing tanks, after which their resulting offspring were behaviorally typed for activity and feeding profiles during the first week of life, following the same procedure used for their mothers (resulting in an additional ∼166 million data points over 9,220 recording hours). This constituted a total of more than 337 million data points obtained over 18,740 hours presented in the present study.

We focus on two main research questions. (i) Does behavioral variation that arises among parents in the apparent absence of genetic and environmental differences predict behavioral differences in the offspring generation? One plausible route linking mother and offspring behavior is via body size: mothers that are more active and/or feed more may become larger and therefore produce larger offspring; larger offspring, in turn, may be more active and/or feed more themselves. Importantly, maternal behavior may also shape offspring size independently of maternal size, for example through differences in resource allocation or provisioning among similarly sized mothers. Thus, we additionally tested whether (ii) the link between mother and offspring behavior (if present) is mediated via maternal size or offspring size.

## RESULTS

### Behavioral individuality in mothers and offspring

As reported previously (14), and consistent with other studies on Amazon mollies (8,12), we find that our 34 mothers exhibit substantial among-individual differences in both behavioral traits: activity (log-transformed, R = 0.356, 95% CI = [0.308, 0.405]) and time spent feeding (R = 0.201, 95% CI = [0.161, 0.246]) - despite the absence of detectable genetic or environmental differences among individuals. These patterns are robust with respect to controlling for individual size and age (adjusted R [95% CI]: log-transformed activity = 0.580 [0.540, 0.627], feeding = 0.275 [0.228, 0.325]). Similarly, we find substantial among-individual differences in activity (log-transformed, R = 0.295, CI = [0.257, 0.337]) and feeding behavior (R = 0.203, CI = [0.174, 0.232]) among their 232 offspring. This variation persists when controlling for offspring size and age as well as mother size at parturition (adjusted R [95% CI]: log-transformed offspring activity = 0.373 [0.323, 0.425], offspring feeding = 0.282 [0.248, 0.319]. See **Supplementary Figure S1** in **Supplementary Information 3** for a visualization of among- and within-individual variation of offspring activity and feeding behavior.

### Maternal behavior predicts offspring activity

We find that mother feeding behavior predicts offspring activity: mothers that spend more time feeding produce offspring that are more active (controlling for offspring feeding behavior) (p = 0.004, partial R^2^ = 0.029, **Figure 1b**; **Supplementary Table S1** in **Supplementary Information 1, Supplementary Figure S2** in **Supplementary Information 3**). We also find a trend suggesting that more active mothers might also produce more active offspring (both log-transformed, p = 0.059, partial R^2^ = 0.009; **Figure 1a**; **Supplementary Table S1** in **Supplementary Information 1**). We note that this trend becomes significant when controlling for mother and offspring size (see below for details). Offspring feeding behavior is not associated with mother activity (log-transformed) (p = 0.593, **Figure 1c**) or mother feeding behavior (p = 0.814, **Figure 1d**) (**Supplementary Table S2** in **Supplementary Information 1**).

**Figure 1.**
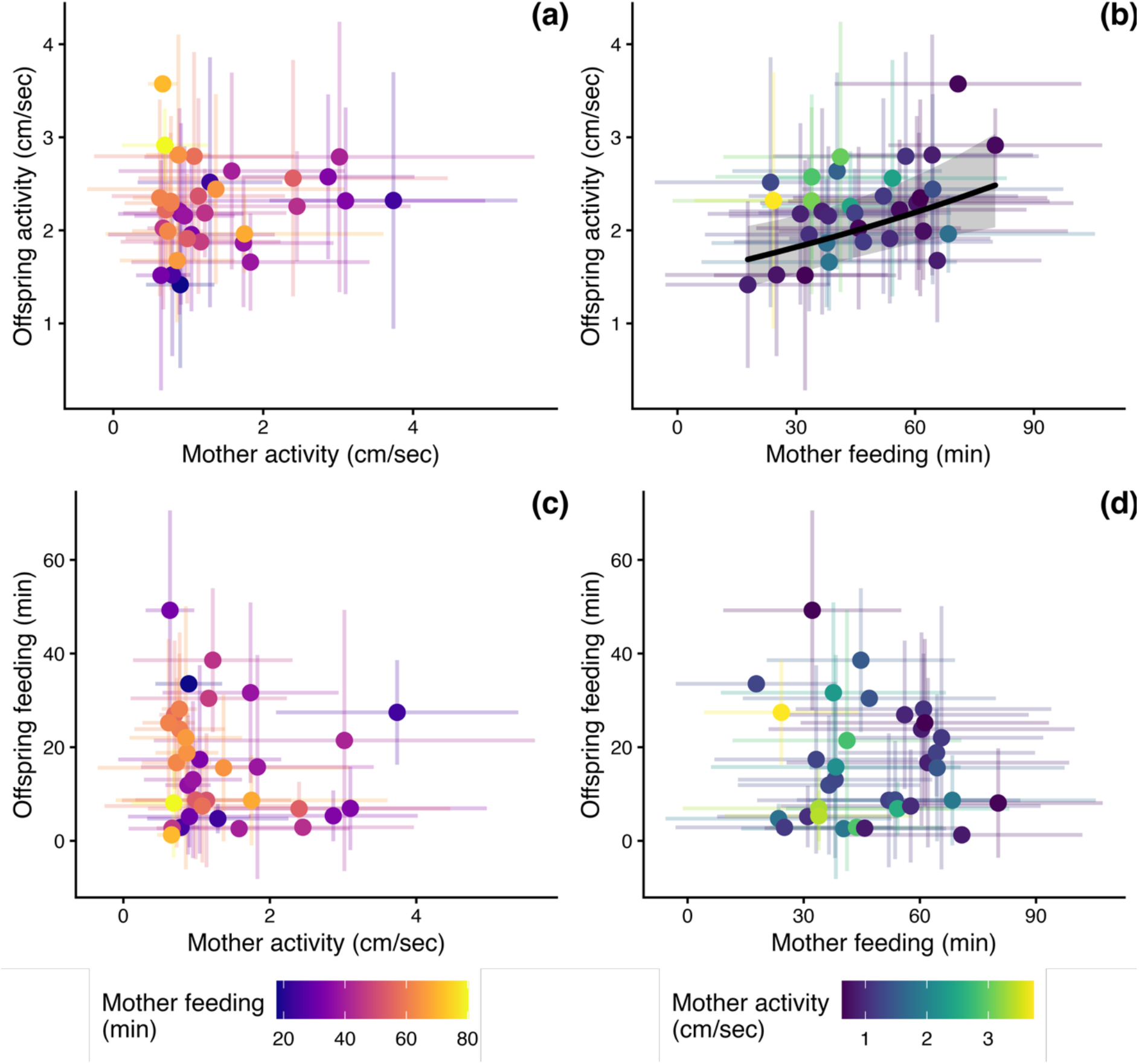
Offspring activity (a) shows a statistically non-significant positive association with mother activity, and (b) is positively correlated with mother feeding. Offspring feeding behavior is not linked to either mother (c) activity or (d) feeding behavior. (a-c) Activity values were log-transformed for statistical analyses and subsequently back-transformed for plotting to aid interpretability. (a-d) Data points represent the mean per mother (N = 34); error bars on the x-axes reflect the standard deviation of daily maternal behavior (based on N_maternal activity_ = 941 and N_maternal feeding_ = 928 data points); error bars on the y-axes reflect the standard deviation in mean offspring behavior (based on N = 922 data points of daily offspring activity and feeding, respectively).

To assess the relationship between maternal and offspring behavior, we used data points for individual offspring (N = 922 daily activity and N = 922 daily feeding measures from N = 232 offspring). An alternative approach (**Supplementary Information 2**) would be to average offspring behavior across all offspring produced by the same mother, yielding a single data point per mother. The slope obtained from such a mid-offspring, mid-parent regression is commonly being used as a measure of narrow-sense heritability in sexually reproducing species (h²) (24–26). When repeating our above analyses at this aggregated level, we find that our above results are robust, i.e., we continue to observe a highly significant link between mother feeding behavior and offspring activity (p = 0.001, partial R^2^ = 0.329), a weaker, non-significant link between mother and offspring activity (p = 0.088, partial R^2^ = 0.097), and no link between mother behavior and offspring feeding (**Supplementary Table S11** in **Supplementary Information 2**). As can be expected, the strength of mother–offspring behavioral associations increases substantially when offspring data are aggregated at the maternal level (thereby collapsing offspring behavioral variation observed among siblings). At this aggregated level, the proportion of variation in offspring activity explained by maternal feeding increases more than tenfold, i.e., the partial R² increases from 2.9% in our offspring-level analysis to 32.9%. Likewise, the explanatory power of maternal activity increases from 0.9% to 9.7% (partial R²).

### Body size does not mediate mother-offspring behavioral associations

In principle, the effects of mother behavior on offspring behavior may be mediated by mother size at parturition or offspring size at birth. However, while offspring size at birth is positively related to offspring activity (log-transformed, p = 0.008, partial R^2^ = 0.011), offspring feeding behavior is not (p = 0.015, partial R^2^ = 0.189; **Supplementary Figure S3a-b** in **Supplementary Information 2; Supplementary Table S4-S5** in **Supplementary Information 1**); and we find no direct link between mother behavior and offspring size at birth (activity p = 0.644; feeding p = 0.623; **Supplementary Figure S3c-d** in **Supplementary Information 2, Supplementary Table 3** in **Supplementary Information 1**). Mother activity (log-transformed, p = 0.227) or feeding (p = 0.664) are not correlated with maternal size at parturition, nor is maternal size at parturition associated with offspring activity (log-transformed, p = 0.536) or feeding (p = 0.267) (**Supplementary Figure S4** in **Supplementary Information 2, Supplementary Tables S6- S8** in **Supplementary Information 1**). We note that, when including both mother size at parturition and offspring size at birth in our main models to test for a mother-offspring behavioral link, the relationship between maternal feeding behavior and offspring activity persists (p = 0.004, partial R^2^ = 0.025), and the non-significant trend for the mother-offspring activity link becomes significant (p = 0.039, partial R^2^ = 0.009) (**Supplementary Tables S9- S10** in **Supplementary Information 1**). This underscores that both mother size at parturition and offspring size at birth do not mediate the link between mother behavior and offspring activity. When repeating these analyses using offspring behavior aggregated at the maternal level (see above), the overall pattern remains unchanged, i.e., we continue to find no evidence that either maternal size at parturition or offspring size at birth mediates the observed mother–offspring behavioral associations (**Supplementary Tables S12- S16** in **Supplementary Information 3**).

## DISCUSSION

Recent work has revealed that substantial, fitness-related phenotypic variation emerges among individuals even in the near absence of genetic and environmental variation. However, whether and to what extent signatures of such seemingly stochastic variation may be transmitted across generations remains a central open question for understanding its potential significance for evolutionary processes. Employing a two-generation behavioral screening with a naturally clonal fish reared individually and in near-identical environments, we found that maternal feeding differences observed during her first 28 days of life predict offspring activity observed during the first days of life. We also found suggestive evidence that maternal activity may positively predict offspring activity, with this relationship becoming statistically significant when controlling for maternal and offspring body size. We found no evidence that body size itself (either maternal size at parturition or offspring size at birth) mediates the observed mother-offspring behavioral associations.

By demonstrating that behavioral differences that arise in the near absence of genetic and environmental differences can be transmitted to the next generation, our results raise the possibility that such stochastically induced phenotypic variation may play an unrecognized role in evolutionary processes. It seems conceivable, for example, that stochastic variation might act as ‘non-genetic standing variation’ that, once selected upon, results in a shift in the distribution of traits in subsequent generations, potentially allowing populations to adaptively track environmental change. While adaptations based on non-genetic sources of variation may be more transient and limited in scope long-term compared to those of a genetic source, non-genetic variation may allow populations to more quickly adapt to rapid environmental change without relying on slower-to-occur genetic mutations (22,27,28). Importantly, while substantial previous research in this area has focused on non-genetic transmission of environmentally induced variation (e.g., transgenerational plasticity (29,30)), our results draw attention to the transmission of variation emerging in the apparent absence of environmental differences, thus broadening the scope and potential importance of non-genetic inheritance in evolution.

Intriguingly, we observe a cross-trait effect (maternal feeding behavior predicts offspring activity) suggesting that stochastically induced behavioral phenotypes may not be inherited as fixed trait values but may instead reflect broader shifts in underlying state variables that manifest across traits. That is, maternal behavior may influence the general physiological or developmental state of offspring, which then shapes behavior indirectly rather than reproducing the same behavioral trait across generations. Importantly, behavioral differences emerging in the apparent absence of genetic and environmental differences may initially emerge through processes such as developmental noise, stochastic variation in gene expression or epigenetic state, minute experiential differences amplified through feedback loops, or causal processes that currently remain unresolved (12–14,31–33). Once established, however, these phenotypic differences may themselves alter maternal physiology, energetic state, endocrine signalling, or epigenetic profiles, thereby creating systematic and predictable pathways through which information about these phenotypes can be transmitted to offspring. While we could not confirm body size (neither maternal size at parturition nor offspring size at birth) as a key mediator linking maternal behavior and offspring activity, other aspects of maternal state may constitute equally likely carriers of information across generations. Variation in such state variables could plausibly persist through reproduction and bias offspring developmental trajectories, even in the absence of detectable differences in body size. Identifying which aspects of maternal state encode and transmit such stochastic variation represents a key challenge for future work.

The strength of mother-offspring behavioral associations varies substantially depending on the level of data aggregation: at the maternal level, where offspring behavior is represented as the mean phenotype of all offspring produced by a given mother, maternal feeding accounts for more than ten times as much variation in offspring activity (32.9%) compared to the offspring-level analysis (2.9%).

A similar, though somewhat less pronounced, pattern emerges for the non-significant trend in maternal activity, whose explanatory power increases from just 0.9% (offspring-level analysis) to 9.7% (maternal-level analysis). This substantial change in effect strength suggests that maternal behavior is associated with a shift in the average behavior of the next generation while, at the same time, individuals vary markedly around this shifted mean. This pattern aligns closely with our previous work showing that even under highly standardized genetic and environmental conditions, Amazon mollies develop substantial behavioral individuality (8,12,14).

Under our experimental conditions, the intergenerational feeding-activity link appears to be transient: maternal feeding behavior predicts offspring activity but there is no reciprocal step, i.e., activity does not predict feeding. The intergenerational link does therefore not seem to extend beyond a single step. This pattern may likely be a result of the highly standardized conditions of our study, in which activity was effectively decoupled from resource acquisition: there was no need to actively search for food as it was provided in a fixed, easily accessible location. Under such conditions, variation in activity carries little functional weight and is therefore unlikely to propagate further across generations. This reduced functional relevance may also explain why the intergenerational activity-activity link was comparatively weak and less consistently detected across analyses. Under more natural environments, where activity directly affects resource acquisition, the observed cross-trait transmission of feeding differences may generate lasting consequences across generations. In such environments, one might also expect a stronger and more persistent activity-activity transmission across generations. Disentangling how environmental factors (particularly those related to resource availability) modulate the functional relevance, persistence, and direction of stochastically induced behavioral phenotypes across multiple generations represents an important direction for future research.

Finally, clonal organisms have been regarded as evolutionary dead ends due to their limited ability to generate genetic diversity. However, our findings suggest that some clonal species may possess a previously underestimated capacity for adaptation; raising the question of whether the Amazon molly, and possibly clonal species more generally, developed mechanisms of enhanced non-genetic inheritance to cope with the loss of genetic diversity. Future studies may address this question by comparing the intergenerational transmission of behavior between asexually reproducing Amazon mollies and closely related sexually reproducing species, such *as P. latipinna* or *P. mexicana*. Upcoming work may also investigate the consequences of the observed non-genetic transmission of behavior for natural Amazon molly populations and their adaptive potential. An exciting experimental approach to test the adaptive potential of Amazon mollies would involve bi-directional selection line experiments. By breeding low-activity and high-activity individuals in distinct lines, for example, and observing their behavioral divergence over multiple generations, we can gain a deeper understanding of the adaptive significance of stochastically induced phenotypic variation in clonal species and its implications for their long-term survival and ecological success.

In conclusion, our findings provide the first empirical evidence that phenotypic variation arising in the near absence of genetic and environmental differences can be transmitted across generations. Our results demonstrate that seemingly stochastic behavioral variation can leave systematic signatures in the next generation, positioning it as biologically meaningful variation with relevance for evolutionary processes. Future work will be essential to identify the underlying mechanisms and to determine how environmental context influences the magnitude and persistence of the observed intergenerational transmission of behavioral phenotypes, thereby clarifying their role in the adaptive potential of natural populations.

## METHODS

### Study animals and holding conditions

Amazon mollies used in our experiment were obtained from a stock population at the Humboldt-Universität zu Berlin, (Berlin, Germany). The lineage used in our experiment was established prior to the start of the experiment by holding a single Amazon molly from the stock population as well as a *P. mexicana* male and their resulting offspring in a 50 liter tank (approx. 20-50 fish, 12:12 hour light:dark cycle, air temperature control at approx. 24±1°C, weekly water changes, fish were fed with Sera vipan baby powder food twice a day). To initiate our experiment, we held potentially gravid females from the stock in individual tanks until they gave birth. This way, we knew exactly when females that we used as mothers in the current experiment were born and could track their mother ID (*N* = 3).

### General procedure

Mothers-to-be were behaviorally phenotyped over the first 37 days of life, comprising the 28-day behavioral typing period analyzed in the present study, followed by an additional nine days of behavioral assays that are analyzed in a separate study (manuscript in preparation; see ‘Behavioral observations’ for details). Following the behavioral typing, females were allowed to reproduce in individual breeding tanks (see ‘Reproduction’). Mother behavioral and reproductive characteristics have been investigated in detail previously (14,16). Once a female produced a brood, her offspring were behaviorally typed in the same manner as their mother - except, the observation period of offspring was shorter compared their mothers’ observation period (mean ± SD: 5.1 ± 2.0 days) in order to process as many offspring as possible with limited tanks space. We note that offspring behavior changed over observation days, with offspring becoming less active and spending more time feeding over time, we therefore controlled offspring observation day throughout (**Supplementary Tables S1- S2, S4-S5, S7-S10** in **Supplementary Information 1**)). For larger broods, we behaviorally typed a random sample of the brood (mean ± SE proportion of offspring sampled per brood = 0.776 ± 0.244 %) with the size of the sample depending on capacity limits (N (behaviorally typed) = 245 offspring, N (total) = 373 offspring).

### Behavioral observations

Behavioral observations for both mothers and their offspring started on the first full day of their life, i.e., individuals were transferred to individual observation tanks (**Supplementary Figure S5** in **Supplementary Information 3**) on the day they were born, and behavioral observations started the next morning. Individuals were recorded from above (Basler acA5472 camera) for 10 hours per day with a resolution of 5 frames per second. During the activity recording (first 8 hours of each day), no food was present in the tank. Every day, individuals were fed for 2 hours after the activity recording in a standardized position (**Supplementary Figure S5** in **Supplementary Information 3**) with a food patch (a mix of Sera vipan baby powder food, agar, and water; for a detailed preparation protocol see Scherer et al. (2023)), which fresh batches being prepared every two to three days.

Activity and feeding recordings were tracked using the software Biotracker (34). Daily activity was assessed as average swimming speed (cm/sec) and time spent feeding (min) was calculated as the amount of time an individual spent in the ‘feeding zone’ (a 5 x 13 cm large zone surrounding the food patch) over the entire feeding period. Quality of tracks was assured via plotting individual movement data (i.e., xy-coordinates over time) in 30 min chunks and visually inspecting the trajectory plots for errors. For the visualization of movement data and the calculation of metrics (daily activity and time spent feeding) we developed a custom-made repository (35).

As part of an auxiliary study that did not bear on the questions addressed in this manuscript (in accordance with 3R guidelines that maximize data collected from individual animals), following the 28-day behavioral observations of mothers, individuals completed a series of standardized behavioral assays, which will be analyzed in detail elsewhere and are briefly described here for completeness and reproducibility (see (36) for a more detailed description of the experimental procedure). Over nine consecutive days, each individual was tested three times in each of three classic paradigms (totaling nine behavioral tests per individual, with one test per days): activity in a novel environment (movement in a new, unfamiliar tank), response to a novel object (mean distance to a newly introduced object), and sociability (time spent in proximity to a transparent container holding two unfamiliar, size-matched conspecifics). All assays were carried out in the same observation tanks used for the 28-day behavioral observations. A detailed description of the procedures is provided in (14). We note that mother activity and feeding behavior did not predict behavior observed during the 9- day behavioral assays, except a positive correlation between early-life activity and new tank activity (**Supplementary Table S17** in **Supplementary Information 4**). Likewise, behavior observed during behavioral assays did not predict offspring activity or feeding behavior (**Supplementary Table S18** in **Supplementary Information 4**).

### Reproduction

Following behavioral observations of the parental generation, females were transferred to individual breeding tanks equipped with ‘Sera biofibres’ (structurally equivalent to thread algae) and a 2 x 4 cm plastic pipe each as refuge (11 liter per tank, flow-through system with approx. 600 liter in total, holding conditions as above). The Amazon molly is a gynogenetic species, i.e., sperm from closely related species is needed to trigger embryonic development but the male’s DNA is not incorporated into the offspring’s genome (37–41). Breeding tanks therefore held one P. mexicana male each. Once a week, we measured females for size using ImageJ (42) and transferred them to a new tank (randomized order) in order to experimentally control for potential male/tank effects. During breeding, females were fed twice a day for 5 days a week with 1/64 tsp (up to the age of 70 days) or 1/32 tsp (from the age of 70 days onwards) of Sea vipan baby powder food. Twice a day (morning and afternoon), we checked for offspring.

We collected each female’s first brood (N = 34 broods). We counted all offspring (mean ± SD brood size = 10.6 ± 5.8 offspring) and measured them for standard length (mean ± SD = 8.1 ± 0.5 mm) (ImageJ, Schneider et al. (2012)) immediately after birth. At parturition, mothers were on average 125.6 ± 18.7 days old (mean ± SD) and 3.7 ± 0.3 cm large (mean ± SD standard length, predicted from individual von Bertalanffy growth curves (43). The above values reflect the final dataset used for statistical analysis. Initially, our sample comprised 287 offspring from 38 mothers. However, we excluded 42 offspring due to missing body size data and an additional 13 offspring for which feeding behavior was not recorded, resulting in a final data set of N = 232 offspring from N = 34 mothers. Accounting for variation in mother size at parturition does not affect our results (**Supplementary Table S9- S10** in **Supplementary Information 1**). Mother age and size at parturition were tightly correlated (linear model with mother size as response and mother age as predictor; N = 34, estimate ± SE = 0.013 ± 0.001, p < 0.001, R^2^ = 0.764), avoiding collinearity, we did not additionally account for variation in mother age at parturition (44).

### Statistical analyses

For analyses, we used R version 4.21 (45) and the following packages: arm (46), data.table (47), dplyr (48), ggplot2 (49), ggpubr (50), lme4 (51), sjPlot (Lüdecke, 2022),tidyverse (53), and viridis (54). For all models, model assumptions were verified using residual and q-q plots. Throughout, mother and offspring activity was log-transformed (natural logarithm) for normality. A complete summary of all full models (containing all predictors) and final models (containing only significant predictors) is provided as **Supplementary Information 1**.

To test whether individuals differed consistently in their behavior from each other, we estimated behavioral repeatabilities. Repeatability is a key parameter to quantify individuality by estimating how much of the total variation observed can be attributed to among-individual differences (55–57). We calculated repeatability for offspring behavior from two linear-mixed effects models with either daily offspring activity or feeding behavior as the response (both models: N = 922 observation days from N = 232 offspring and N = 34 mothers). Offspring ID, mother ID, grandmother ID, and tank ID were included as random terms (no predictors were included, aka null model) (following Hertel et al. (2020). Additionally, we adjusted offspring repeatabilities for individual size at birth and observation day by including these variables as predictors in the respective null model. Similarly, repeatability of mother behavior was assessed from two linear mixed effects models with either mother daily activity (N = 941 observations from 34 individuals) or mother daily time spent feeding (N = 928 observations from 34 individuals) as response, and grandmother ID (N = 3 females our experimental mothers originate from) as predictor. Behavioral repeatabilities for mothers were adjusted for individual size and age by including their predicted size of the week and age in weeks (as a categorical variation as predictor variables in the respective null model. We also included the size-age interaction term as predictor to account for the fact that the effect of size on behavior varied among observation weeks. Mother repeatabilities were not controlled for tank ID as each tank was only used twice in the maternal generation.

To test whether maternal behavior predicts offspring behavior, we fitted two complementary linear mixed-effects models. In the first model, offspring activity was modeled as a function of maternal activity and maternal feeding behavior, with grandmother ID and offspring observation day included as control variables. Offspring ID and mother ID, as well offspring tank specifics were included as random term. Tank specifics included: tank ID (N = 24), tank system (N = 4), tank distance to filter (1m, 2m, 3m), and tank centrality (central vs. peripheral, with peripheral referring to a tank’s position within a filter system). In a parallel model, we tested whether maternal behavior predicted brood-level offspring feeding behavior, using offspring feeding as the response and the same predictor structure. For both models, we included daily behavioral measures, yielding several data points per offspring and behavior (activity or feeding), resulting in N = 922 total observations from N = 232 offspring and N = 34 mothers, respectively. Activity and feeding behavior were moderately negatively correlated in both mothers (p = 0.014, R^2^ = 0.175) and offspring (p = 0.011, R^2^ = 0.185) (linear model with average feeding as predictor and average activity as response, N = 34 in both models). Including both traits simultaneously in each model therefore allowed us to estimate their independent effects while controlling for covariation between behaviors, rather than treating them as independent.

To test whether offspring size plays a mediating role in linking mother and offspring behavior, we first tested whether mother behavior predicts offspring size at birth by building a linear mixed-effects model with offspring size as response and average maternal activity as well as average maternal time spent feeding as predictors (N = 232, i.e., one data point per offspring). Grandmother ID was included as a control factor and mother ID was included as random term. We then tested whether offspring size predicts offspring behavior (activity or feeding) by fitting two linear mixed-effects models, one with daily offspring activity and another one with daily offspring feeding as response. In both models, offspring size was included as the main predictor, grandmother ID as a control factor, and mother ID as well as offspring tank specifics (tank ID, tank system, distance to the filter, and tank centrality) were included as random terms (N = 922 total observations from N = 232 offspring and N = 34 mothers). In parallel, we tested for an alternative mediation pathway via mother size using the same model structure as before. That is, we first built a linear model with maternal size at parturition as response, and maternal activity and feeding behavior as predictors (N = 34 broods, grandmother ID was included as a control factor). We then fitted two linear mixed-effects models with either offspring activity or feeding behavior as response, mother size at parturition as predictor, and the same control structure as before (i.e., grandmother ID was included as a factor and as random term, we included mother ID, tank ID, tank system, distance to the filter, and tank centrality (N = 922 total observations from N = 232 offspring and N = 34 mothers).

### Use of AI-assisted tools

The AI-based language tool ChatGPT (version 5.3) was used to assist with language editing and stylistic refinement. All scientific content, including data analysis, and conclusions were developed by the authors.

### Data availability statement

The datasets supporting this article have been uploaded on Figshare (59).

