## Supplementary Information for "Non-genetic inheritance of stochastically induced behavioral individuality in a naturally clonal fish"

### Supplementary Information 1: Model summaries main analyses

**Supplementary Table 1**. Results of a linear mixed-effects model testing whether mother behavior (activity, feeding) predicts offspring activity. Daily offspring activity measures are used, with observation day being included as a covariate to account for variation in offspring behavior caused by offspring age. The result is controlled for both grandmother ID (included as a fixed effect, N = 3) and mother ID (included as a random effect, N = 34). To control for variation among offspring potentially arising through tank differences, the model includes the following variables as random effects: tank ID (N = 24), tank system (N = 4), tank distance to filter (1m, 2m, 3m), tank centrality (central vs. peripheral).

| Response | Predictors | Estimates | SE | Statistic | p | df | partial $R^2$ |
|---|---|---|---|---|---|---|---|
| Offspring activity (cm/sec, log-transformed) | (Intercept) | 0.765 | 0.125 | - | - | - | - |
| | Mother activity (cm/sec, log-transformed) | 0.114 | 0.058 | 3.574 | 0.059 | 1 | 0.009 |
| | Mother feeding (min) | 0.006 | 0.002 | 8.255 | 0.004 | 1 | 0.029 |
| | Grandmother ID [m2] | 0.031 | 0.083 | 3.689 | 0.158 | 2 | 0.012 |
| | Grandmother ID [m3] | -0.086 | 0.056 | | | | |
| | Observation day | -0.117 | 0.006 | 278.59 | <0.001 | 1 | 0.116 |
| | **Random Effects** | | | | | | |
| | $\sigma^2$ | 0.09 | | | | | |
| | $\tau_{00}$ Offspring ID | 0.11 | | | | | |
| | $\tau_{00}$ Mother ID | 0.00 | | | | | |
| | $\tau_{00}$ Offspring tank ID | 0.09 | | | | | |
| | $\tau_{00}$ Offspring tank system | 0.00 | | | | | |
| | $\tau_{00}$ Offspring tank distance to filter | 0.00 | | | | | |
| | $\tau_{00}$ Offspring tank centrality | 0.00 | | | | | |
| | ICC | 0.69 | | | | | |
| | N Offspring ID | 232 | | | | | |
| | N Mother ID | 34 | | | | | |
| | N Offspring tank ID | 24 | | | | | |
| | N Offspring tank system | 4 | | | | | |
| | N Offspring tank distance to filter | 3 | | | | | |
| | N Offspring tank centrality | 2 | | | | | |
| | Observations | 922 | | | | | |
| | Marginal $R^2$ / Conditional $R^2$ | 0.154 / 0.739 | | | | | |
| | AIC | 2065.456 | | | | | |

**Supplementary Table 2**. Results of a linear mixed-effects model testing whether mother behavior (activity, feeding) predicts offspring feeding. Daily offspring feeding measures are used, with observation day being included as a covariate as to account for variation in offspring behavior caused by offspring age. The result is controlled for both grandmother ID (included as a fixed effect, N = 3) and mother ID (included as a random effect, N = 34). To control for variation among offspring potentially arising through tank differences, the model includes the following variables as random effects: tank ID (N = 24), tank system (N = 4), tank distance to filter (1m, 2m, 3m), tank centrality (central vs. peripheral).

| Response | Predictors | Estimates | SE | $\chi^2$ | p | df | partial $R^2$ |
|---|---|---|---|---|---|---|---|
| Offspring feeding (min) | (Intercept) | 4.293 | 6.482 | - | - | - | - |
| | Mother feeding (min) | 0.065 | 0.118 | 0.055 | 0.814 | 1 | 0.001 |
| | Mother activity (cm/sec, log-transformed) | 0.885 | 3.750 | 0.286 | 0.593 | 1 | 0.002 |
| | Grandmother ID [m2] | -1.469 | 5.180 | 6.780 | 0.034 | 2 | 0.032 |
| | Grandmother ID [m4] | -9.103 | 3.456 | | | | |
| | Observation day | 5.167 | 0.352 | 189.160 | <0.001 | 1 | 0.120 |
| | **Random Effects** | | | | | | |
| | $\sigma^2$ | 265.90 | | | | | |
| | $\tau_{00}$ Offspring ID | 130.41 | | | | | |
| | $\tau_{00}$ Mother ID | 41.53 | | | | | |
| | $\tau_{00}$ Offspring tank ID | 17.91 | | | | | |
| | $\tau_{00}$ Offspring tank system | 0.00 | | | | | |
| | $\tau_{00}$ Offspring tank distance to filter | 0.00 | | | | | |
| | $\tau_{00}$ Offspring tank centrality | 6.64 | | | | | |
| | N Offspring ID | 232 | | | | | |
| | N Mother ID | 34 | | | | | |
| | N Offspring tank ID | 24 | | | | | |
| | N Offspring tank system | 4 | | | | | |
| | N Offspring tank distance to filter | 3 | | | | | |
| | N Offspring tank centrality | 2 | | | | | |
| | Observations | 922 | | | | | |
| | Marginal $R^2$ / Conditional $R^2$ | 0.264 / NA | | | | | |
| | AIC | 8054.281 | | | | | |

**Supplementary Table 3**: Result of a linear mixed-effects model testing whether maternal behavior (activity, feeding) predicts offspring size at birth. The result is controlled for both grandmother ID (N = 3) and mother ID (included as a random term, N = 34).

| Response | Predictors | Estimates | SE | $\chi^2$ | p | df | partial $R^2$ |
|---|---|---|---|---|---|---|---|
| Offspring size (mm) | (Intercept) | 8.117 | 0.240 | - | - | - | - |
| | Mother activity (cm/sec, log-transformed) | -0.073 | 0.158 | 0.213 | 0.644 | 1 | 0.002 |
| | Mother feeding (min) | 0.002 | 0.005 | 0.242 | 0.623 | 1 | 0.005 |
| | Grandmother ID [m2] | 0.275 | 0.217 | 14.338 | 0.001 | 2 | 0.221 |
| | Grandmother ID [m3] | -0.453 | 0.145 | | | | |
| | **Random Effects** | | | | | | |
| | $\sigma^2$ | 0.16 | | | | | |
| | $\tau_{00}$ Mother ID | 0.11 | | | | | |
| | ICC | 0.41 | | | | | |
| | N Mother ID | 34 | | | | | |
| | Observations | 232 | | | | | |
| | Marginal $R^2$ / Conditional $R^2$ | 0.237 / 0.551 | | | | | |
| | AIC | 314.451 | | | | | |

**Supplementary Table 4**. Results of a linear mixed-effects model testing whether offspring size at birth predicts offspring activity. Daily offspring activity measures are used, with observation day being included as a covariate as to account for variation in offspring behavior caused by offspring age. The result is controlled for both grandmother ID (included as a fixed effect, N = 3) and mother ID (included as a random effect, N = 34). To control for variation among offspring potentially arising through tank differences, the model includes the following variables as random effects: tank ID (N = 24), tank system (N = 4), tank distance to filter (1m, 2m, 3m), tank centrality (central vs. peripheral).

| Response | Predictors | Estimates | SE | $\chi^2$ | p | df | partial $R^2$ |
|---|---|---|---|---|---|---|---|
| Offspring activity (cm/sec, log-transformed) | (Intercept) | -0.057 | 0.406 | - | - | - | - |
| | Offspring size (mm) | 0.136 | 0.049 | 6.981 | 0.008 | 1 | 0.011 |
| | Grandmother ID [m2] | -0.028 | 0.081 | 0.208 | 0.901 | 2 | 0.000 |
| | Grandmother ID [m4] | -0.024 | 0.060 | | | | |
| | Observation day | -0.119 | 0.006 | 283.76 | <0.001 | 1 | 0.120 |
| | **Random Effects** | | | | | | |
| | $\sigma^2$ | 0.09 | | | | | |
| | $\tau_{00}$ Offspring ID | 0.10 | | | | | |
| | $\tau_{00}$ Mother ID | 0.00 | | | | | |
| | $\tau_{00}$ Offspring tank ID | 0.09 | | | | | |
| | $\tau_{00}$ Offspring tank system | 0.00 | | | | | |
| | $\tau_{00}$ Offspring tank distance to filter | 0.00 | | | | | |
| | $\tau_{00}$ Offspring tank centrality | 0.00 | | | | | |
| | N Offspring ID | 232 | | | | | |
| | N Mother ID | 34 | | | | | |
| | N Offspring tank ID | 24 | | | | | |
| | N Offspring tank system | 4 | | | | | |
| | N Offspring tank distance to filter | 3 | | | | | |
| | N Offspring tank centrality | 2 | | | | | |
| | Observations | 922 | | | | | |
| | Marginal $R^2$ / Conditional $R^2$ | 0.341 / NA | | | | | |
| | AIC | 2054.345 | | | | | |

**Supplementary Table 5**. Results of a linear mixed-effects model testing whether offspring size at birth predicts offspring feeding. Daily offspring feeding measures are used, with observation day being included as a covariate as to account for variation in offspring behavior caused by offspring age. . The result is controlled for both grandmother ID (included as a fixed effect, N = 3) and mother ID (included as a random effect, N = 34). To control for variation among offspring potentially arising through tank differences, the model includes the following variables as random effects: tank ID (N = 24), tank system (N = 4), tank distance to filter (1m, 2m, 3m), tank centrality (central vs. peripheral).

| Response | Predictors | Estimates | SE | $\chi^2$ | p | df | partial $R^2$ |
|---|---|---|---|---|---|---|---|
| Offspring feeding (min) | (Intercept) | -13.826 | 18.404 | - | - | - | - |
| | Offspring size (mm) | 2.575 | 2.204 | 1.336 | 0.248 | 1 | 0.005 |
| | Grandmother ID [m2] | -2.391 | 4.814 | 4.851 | 0.089 | 2 | 0.023 |
| | Grandmother ID [m4] | -7.807 | 3.446 | | | | |
| | Observation day | 5.148 | 0.351 | 188.580 | <0.001 | 1 | 0.122 |
| | **Random Effects** | | | | | | |
| | $\sigma^2$ | 265.77 | | | | | |
| | $\tau_{00}$ Offspring ID | 130.29 | | | | | |
| | $\tau_{00}$ Mother ID | 40.72 | | | | | |
| | $\tau_{00}$ Offspring tank ID | 17.34 | | | | | |
| | $\tau_{00}$ Offspring tank system | 0.00 | | | | | |
| | $\tau_{00}$ Offspring tank position | 0.00 | | | | | |
| | $\tau_{00}$ Offspring tank centrality | 6.98 | | | | | |
| | N Offspring ID | 232 | | | | | |
| | N Mother ID | 34 | | | | | |
| | N Offspring tank ID | 24 | | | | | |
| | N Offspring tank system | 4 | | | | | |
| | N Offspring tank position | 3 | | | | | |
| | N Offspring tank centrality | 2 | | | | | |
| | Observations | 922 | | | | | |
| | Marginal $R^2$ / Conditional $R^2$ | 0.268 / NA | | | | | |
| | AIC | 8.049.681 | | | | | |

**Supplementary Table 6**. Results of a linear model testing whether maternal behavior (activity, feeding) predicts mother size at parturition. The result is controlled for grandmother ID, included as a fixed effect because the number of grandmothers is too low to be included as a random term, N = 3).

<table>
<tr><th>Response</th><th>Predictors</th><th>Estimates</th><th>SE</th><th>F</th><th>p</th><th>df</th><th>partial $R^2$</th></tr>
<tr><td rowspan="8">Mother size at parturition (cm)</td><td>(Intercept)</td><td>3.760</td><td>0.099</td><td>-</td><td>-</td><td>-</td><td>-</td></tr>
<tr><td>Mother activity<br>(cm/sec, log-transformed)</td><td>0.077</td><td>0.062</td><td>1.521</td><td>0.227</td><td>1</td><td>0.051</td></tr>
<tr><td>Mother feeding (min)</td><td>0.001</td><td>0.002</td><td>0.1922</td><td>0.664</td><td>1</td><td>0.007</td></tr>
<tr><td>Grandmother ID [m2]</td><td>0.244</td><td>0.098</td><td rowspan="2">35.999</td><td rowspan="2"><0.001</td><td rowspan="2">2</td><td rowspan="2">0.713</td></tr>
<tr><td>Grandmother ID [m3]</td><td>-0.397</td><td>0.062</td></tr>
<tr><td>Observations</td><td colspan="6">34</td></tr>
<tr><td>$R^2$ / $R^2$ adjusted</td><td colspan="6">0.723 / 0.684</td></tr>
<tr><td>AIC</td><td colspan="6">-21.904</td></tr>
</table>

**Supplementary Table 7**. Results of a linear mixed-effects model testing whether mother size at parturition predicts offspring activity. Daily offspring activity measures are used, with observation day being included as a covariate as to account for variation in offspring behavior caused by offspring age. The result is controlled for both grandmother ID (included as a fixed effect, N = 3) and mother ID (included as a random effect, N = 34). To control for variation among offspring potentially arising through tank differences, the model includes the following variables as random effects: tank ID (N = 24), tank system (N = 4), tank distance to filter (1m, 2m, 3m), tank centrality (central vs. peripheral).

| Response | Predictors | Estimates | SE | Statistic | p | df | partial R2 |
|---|---|---|---|---|---|---|---|
| Offspring activity (cm/sec, log-transformed) | (Intercept) | 0.544 | 0.830 | - | - | - | - |
| | Mother size at parturition (cm) | 0.135 | 0.218 | 0.383 | 0.536 | 1 | 0.000 |
| | Grandmother ID [m2] | -0.027 | 0.111 | 0.1828 | 0.913 | 2 | 0.000 |
| | Grandmother ID [m4] | -0.028 | 0.097 | | | | |
| | Observation day | -0.118 | 0.006 | 280.38 | <0.001 | 1 | 0.118 |
| | **Random Effects** | | | | | | |
| | $\sigma^2$ | 0.09 | | | | | |
| | $\tau_{00}$ Offspring ID | 0.11 | | | | | |
| | $\tau_{00}$ Mother ID | 0.01 | | | | | |
| | $\tau_{00}$ Offspring tank ID | 0.09 | | | | | |
| | $\tau_{00}$ Offspring tank system | 0.00 | | | | | |
| | $\tau_{00}$ Offspring tank distance to filter | 0.00 | | | | | |
| | $\tau_{00}$ Offspring tank centrality | 0.00 | | | | | |
| | N Offspring ID | 232 | | | | | |
| | N Mother ID | 34 | | | | | |
| | N Offspring tank ID | 24 | | | | | |
| | N Offspring tank system | 4 | | | | | |
| | N Offspring tank distance to filter | 3 | | | | | |
| | N Offspring tank centrality | 2 | | | | | |
| | Observations | 922 | | | | | |
| | Marginal $R^2$ / Conditional $R^2$ | 0.320 / NA | | | | | |
| | AIC | 2057.434 | | | | | |

**Supplementary Table 8**. Results of a linear mixed-effects model testing whether mother size at parturition predicts offspring feeding. Daily offspring feeding measures are used, with observation day being included as a covariate as to account for variation in offspring behavior caused by offspring age. The result is controlled for both grandmother ID (included as a fixed effect, N = 3) and mother ID (included as a random effect, N = 34). To control for variation among offspring potentially arising through tank differences, the model includes the following variables as random effects: tank ID (N = 24), tank system (N = 4), tank distance to filter (1m, 2m, 3m), tank centrality (central vs. peripheral).

| Response | Predictors | Estimates | SE | $\chi^2$ | p | df | partial $R^2$ |
|---|---|---|---|---|---|---|---|
| Offspring feeding (min) | (Intercept) | 53.348 | 41.375 | - | - | - | - |
| | Mother size at parturition (cm) | -12.142 | 10.872 | 1.234 | 0.267 | 1 | 0.004 |
| | Grandmother ID [m2] | 1.886 | 5.791 | 6.277 | 0.043 | 2 | 0.025 |
| | Grandmother ID [m4] | -13.126 | 5.038 | | | | |
| | Observation day | 5.178 | 0.352 | 190.630 | <0.001 | 1 | 0.118 |
| | **Random Effects** | | | | | | |
| | $\sigma^2$ | 265.96 | | | | | |
| | $\tau_{00}$ Offspring ID | 128.87 | | | | | |
| | $\tau_{00}$ Mother ID | 41.70 | | | | | |
| | $\tau_{00}$ Offspring tank ID | 18.42 | | | | | |
| | $\tau_{00}$ Offspring tank system | 0.00 | | | | | |
| | $\tau_{00}$ Offspring tank filter distance | 0.00 | | | | | |
| | $\tau_{00}$ Offspring tank centrality | 6.65 | | | | | |
| | N Offspring ID | 232 | | | | | |
| | N Mother ID | 34 | | | | | |
| | N Offspring tank ID | 24 | | | | | |
| | N Offspring tank system | 4 | | | | | |
| | N Offspring tank position | 3 | | | | | |
| | N Offspring tank centrality | 2 | | | | | |
| | Observations | 922 | | | | | |
| | Marginal $R^2$ / Conditional $R^2$ | 0.267 / NA | | | | | |
| | AIC | 8046.581 | | | | | |

**Supplementary Table 9**. Results of a linear mixed-effects model testing whether mother behavior (activity, feeding) predicts offspring activity. The model structure is identical to the model presented in Supplementary Table 1, except that maternal size at parturition and offspring size at birth are included as additional covariates.

| Response | Predictors | Estimates | SE | $\chi^2$ | p | df | partial $R^2$ |
|---|---|---|---|---|---|---|---|
| Offspring activity (cm/sec, log-transformed) | (Intercept) | -0.212 | 0.776 | - | - | - | - |
| | Mother activity (cm/sec, log-transformed) | 0.121 | 0.058 | 4.243 | 0.039 | 1 | 0.009 |
| | Mother feeding (min) | 0.006 | 0.002 | 8.425 | 0.004 | 1 | 0.025 |
| | Grandmother ID [m2] | -0.016 | 0.095 | 0.394 | 0.821 | 2 | 0.012 |
| | Grandmother ID [m4] | -0.049 | 0.086 | | | | |
| | Offspring size at birth (mm) | 0.135 | 0.049 | 7.316 | 0.007 | 1 | 0.003 |
| | Mother size at parturition (cm) | -0.027 | 0.205 | 0.017 | 0.897 | 1 | 0.115 |
| | Observation day | -0.118 | 0.006 | 282.72 | <0.001 | 1 | 0.027 |
| | **Random Effects** | | | | | | |
| | $\sigma^2$ | 0.09 | | | | | |
| | $\tau_{00}$ Offspring ID | 0.10 | | | | | |
| | $\tau_{00}$ Mother ID | 0.00 | | | | | |
| | $\tau_{00}$ Offspring tank ID | 0.09 | | | | | |
| | $\tau_{00}$ Offspring tank system | 0.00 | | | | | |
| | $\tau_{00}$ Offspring tank position | 0.00 | | | | | |
| | $\tau_{00}$ Offspring tank centrality | 0.00 | | | | | |
| | N Offspring ID | 232 | | | | | |
| | N Mother ID | 34 | | | | | |
| | N Offspring tank ID | 24 | | | | | |
| | N Offspring tank system | 4 | | | | | |
| | N Offspring tank position | 3 | | | | | |
| | N Offspring tank centrality | 2 | | | | | |
| | Observations | 922 | | | | | |
| | Marginal $R^2$ / Conditional $R^2$ | 0.384 / NA | | | | | |
| | AIC | 2067.373 | | | | | |

**Supplementary Table 10**. Results of a linear mixed-effects model testing whether mother behavior (activity, feeding) predicts offspring feeding. The model structure is identical to the model presented in Supplementary Table 2, except that maternal size at parturition and offspring size at birth are included as additional covariates.

| *Response* | *Predictors* | *Estimates* | *SE* | *Statistic* | *p* | *df* | *partial R2* |
|---|---|---|---|---|---|---|---|
| Offspring feeding (min) | (Intercept) | 41.193 | 42.143 | - | - | - | - |
| | Mother feeding (min) | 0.056 | 0.112 | 0.242 | 0.623 | 1 | 0.006 |
| | Mother activity (cm/sec, log-transformed) | 2.510 | 3.654 | 1.162 | 0.497 | 1 | 0.002 |
| | Grandmother ID [m2] | 1.826 | 5.608 | 0.461 | 0.034 | 2 | 0.031 |
| | Grandmother ID [m4] | -13.952 | 5.096 | | | | |
| | Offspring size at birth (mm) | 3.554 | 2.264 | 2.351 | 0.125 | 1 | 0.007 |
| | Mother size at parturition (cm) | -17.232 | 11.131 | 2.329 | 0.127 | 1 | 0.009 |
| | Observation day | 5.176 | 0.352 | 282.72 | <0.001 | 1 | 0.126 |
| | **Random Effects** | | | | | | |
| | $\sigma^2$ | 265.72 | | | | | |
| | $\tau_{00}$ Offspring ID | 131.30 | | | | | |
| | $\tau_{00}$ Mother ID | 34.11 | | | | | |
| | $\tau_{00}$ Offspring tank ID | 16.60 | | | | | |
| | $\tau_{00}$ Offspring tank system | 0.00 | | | | | |
| | $\tau_{00}$ Offspring tank position | 0.00 | | | | | |
| | $\tau_{00}$ Offspring tank centrality | 6.42 | | | | | |
| | N Offspring ID | 232 | | | | | |
| | N Mother ID | 34 | | | | | |
| | N Offspring tank ID | 24 | | | | | |
| | N Offspring tank system | 4 | | | | | |
| | N Offspring tank position | 3 | | | | | |
| | N Offspring tank centrality | 2 | | | | | |
| | Observations | 922 | | | | | |
| | Marginal $R^2$ / Conditional $R^2$ | 0.280 / NA | | | | | |
| | AIC | 8045.051 | | | | | |

### Supplementary Information 2: Robustness towards data aggregation

**Supplementary Table 11.** Results of linear models testing whether maternal behavior (activity, feeding) predicts offspring activity (top) or feeding behavior (bottom) at the brood level. The results are controlled for grandmother ID (included as a fixed effect, N = 3).

| Response | Predictors | Estimates | SE | F | p | df | partial $R^2$ |
|---|---|---|---|---|---|---|---|
| Offspring activity (cm/sec, log-transformed; brood average) | (Intercept) | 0.388 | 0.106 | - | - | - | - |
| | Mother activity (cm/sec, log-transformed) | 0.118 | 0.067 | 3.111 | 0.088 | 1 | 0.097 |
| | Mother feeding (min) | 0.008 | 0.002 | 14.215 | 0.001 | 1 | 0.329 |
| | Grandmother ID [m2] | 0.157 | 0.105 | 1.576 | 0.224 | 2 | 0.098 |
| | Grandmother ID [m3] | -0.018 | 0.066 | | | | |
| | Observations | 34 | | | | | |
| | $R^2$ / $R^2$ adjusted | 0.377 / 0.291 | | | | | |
| | AIC | 36.397 | | | | | |
| Offspring feeding (min, brood average) | (Intercept) | 26.212 | 6.793 | - | - | - | - |
| | Mother feeding (min) | -0.065 | 0.133 | 0.240 | 0.628 | 1 | 0.008 |
| | Mother activity (cm/sec, log-transformed) | -1.429 | 4.261 | 0.112 | 0.741 | 1 | 0.004 |
| | Grandmother ID [m2] | -7.470 | 6.704 | 4.310 | 0.023 | 2 | 0.229 |
| | Grandmother ID [m3] | -12.401 | 4.224 | | | | |
| | Observations | 34 | | | | | |
| | $R^2$ / $R^2$ adjusted | 0.273 / 0.172 | | | | | |
| | AIC | 265.649 | | | | | |

**Supplementary Table 12:** Results of a linear model testing whether maternal behavior (activity, feeding) predicts offspring size at birth. The result is controlled for grandmother ID (included as a fixed effect because the number of grandmothers is too low to be included as a random term, N = 3).

<table>
<tr><th>Response</th><th>Predictors</th><th>Estimates</th><th>SE</th><th>F</th><th>p</th><th>df</th><th>partial $R^2$</th></tr>
<tr><td rowspan="8">Offspring size<br>(mm, brood average)</td><td>(Intercept)</td><td>8.099</td><td>0.259</td><td>-</td><td>-</td><td>-</td><td>-</td></tr>
<tr><td>Mother activity<br>(cm/sec, log-transformed)</td><td>-0.088</td><td>0.162</td><td>0.290</td><td>0.594</td><td>1</td><td>0.010</td></tr>
<tr><td>Mother feeding (min)</td><td>0.003</td><td>0.005</td><td>0.474</td><td>0.497</td><td>1</td><td>0.016</td></tr>
<tr><td>Grandmother ID [m2]</td><td>0.276</td><td>0.256</td><td rowspan="2">7.337</td><td rowspan="2">0.003</td><td rowspan="2">2</td><td rowspan="2">0.336</td></tr>
<tr><td>Grandmother ID [m3]</td><td>-0.474</td><td>0.161</td></tr>
<tr><td>Observations</td><td colspan="6">34</td></tr>
<tr><td>$R^2$ / $R^2$ adjusted</td><td colspan="6">0.349 / 0.259</td></tr>
<tr><td>AIC</td><td colspan="6">43.490</td></tr>
</table>

**Supplementary Table 13**. Results of a linear model testing whether offspring size at birth predicts offspring activity (top) or offspring feeding (bottom) at the brood level. The results are controlled for grandmother ID (included as a fixed effect because the number of grandmothers is too low to be included as a random term, N = 3).

| Response | Predictors | Estimates | SE | F | p | df | partial $R^2$ |
|---|---|---|---|---|---|---|---|
| Offspring activity (cm/sec, log-transformed; brood average) | (Intercept) | -0.417 | 0.638 | - | - | - | - |
| | Offspring size (mm, brood average) | 0.171 | 0.079 | 4.672 | 0.039 | 1 | 0.139 |
| | Offspring feeding (min, brood average) | -0.011 | 0.003 | 12.718 | 0.002 | 1 | 0.295 |
| | Grandmother ID [m2] | 0.053 | 0.103 | 12.110 | 0.814 | 2 | 0.014 |
| | Grandmother ID [m3] | -0.017 | 0.078 | | | | |
| | Observations | 34 | | | | | |
| | $R^2$ / $R^2$ adjusted | 0.363 / 0.275 | | | | | |
| | AIC | 37.142 | | | | | |
| Offspring feeding (min, brood average) | (Intercept) | -40.078 | 31.881 | - | - | - | - |
| | Offspring size (mm, brood average) | 10.165 | 3.911 | 6.754 | 0.015 | 1 | 0.189 |
| | Offspring activity (cm/sec, log-transformed; brood average) | -27.647 | 7.945 | 12.110 | 0.002 | 1 | 0.295 |
| | Grandmother ID [m2] | -5.122 | 5.186 | 1.853 | 0.175 | 2 | 0.113 |
| | Grandmother ID [m3] | -6.984 | 3.781 | | | | |
| | Observations | 34 | | | | | |
| | $R^2$ / $R^2$ adjusted | 0.528 / 0.463 | | | | | |
| | AIC | 250.933 | | | | | |

**Supplementary Table 14**. Results of a linear model testing whether maternal behavior (activity, feeding) predicts mother size at parturition. The result is controlled for grandmother ID (included as a fixed effect because the number of grandmothers is too low to be included as a random term, N = 3).

<table>
<tr><th>Response</th><th>Predictors</th><th>Estimates</th><th>SE</th><th>F</th><th>p</th><th>df</th><th>partial R²</th></tr>
<tr><td rowspan="8">Mother size at parturition (cm)</td><td>(Intercept)</td><td>3.760</td><td>0.099</td><td>-</td><td>-</td><td>-</td><td>-</td></tr>
<tr><td>Mother activity<br>(cm/sec, log-transformed)</td><td>0.077</td><td>0.062</td><td>1.521</td><td>0.227</td><td>1</td><td>0.051</td></tr>
<tr><td>Mother feeding (min)</td><td>0.001</td><td>0.002</td><td>0.1922</td><td>0.664</td><td>1</td><td>0.007</td></tr>
<tr><td>Grandmother ID [m2]</td><td>0.244</td><td>0.098</td><td rowspan="2">35.999</td><td rowspan="2"><0.001</td><td rowspan="2">2</td><td rowspan="2">0.713</td></tr>
<tr><td>Grandmother ID [m3]</td><td>-0.397</td><td>0.062</td></tr>
<tr><td>Observations</td><td colspan="6">34</td></tr>
<tr><td>$R^2$ / $R^2$ adjusted</td><td colspan="6">0.723 / 0.684</td></tr>
<tr><td>AIC</td><td colspan="6">-21.904</td></tr>
</table>

**Supplementary Table 15.** Results of a linear model testing whether maternal size at parturition predicts offspring activity (top) or offspring feeding (bottom) at the brood level. The results are controlled for grandmother ID (N = 3).

| Response | Predictors | Estimates | SE | F | p | df | partial $R^2$ |
|---|---|---|---|---|---|---|---|
| Offspring activity (cm/sec, log-transformed; brood average) | (Intercept) | 0.714 | 0.820 | - | - | - | - |
| | Mother size at parturition (cm) | 0.062 | 0.213 | 0.084 | 0.074 | 1 | 0.003 |
| | Offspring feeding (min, brood average) | -0.009 | 0.003 | 7.358 | 0.011 | 1 | 0.202 |
| | Grandmother ID [m2] | 0.085 | 0.123 | 0.306 | 0.739 | 2 | 0.021 |
| | Grandmother ID [m3] | -0.050 | 0.116 | | | | |
| | Observations | 34 | | | | | |
| | $R^2$ / $R^2$ adjusted | 0.262 / 0.160 | | | | | |
| | AIC | 42.123 | | | | | |
| Offspring feeding (min, brood average) | (Intercept) | 57.617 | 42.205 | - | - | - | - |
| | Mother size at parturition (cm) | -4.406 | 11.133 | 0.157 | 0.695 | 1 | 0.005 |
| | Offspring activity (cm/sec, log-transformed; brood average) | -23.584 | 8.694 | 7.358 | 0.011 | 1 | 0.202 |
| | Grandmother ID [m2] | -2.598 | 6.473 | 3.419 | 0.046 | 2 | 0.191 |
| | Grandmother ID [m3] | -13.738 | 5.514 | | | | |
| | Observations | 34 | | | | | |
| | $R^2$ / $R^2$ adjusted | 0.421 / 0.342 | | | | | |
| | AIC | 257.869 | | | | | |

**Supplementary Table 16:** Results of linear models testing whether maternal behavior (activity, feeding) predicts offspring activity (top) or feeding behavior (bottom) at the brood level. The model structure is identical to the models presented in Supplementary Table 10, except that maternal size at parturition and offspring size at birth are included as additional covariates.

| Response | Predictors | Estimates | SE | F | p | df | partial $R^2$ |
|---|---|---|---|---|---|---|---|
| Offspring activity (cm/sec, log-transformed; brood average) | (Intercept) | -0.023 | 0.828 | - | - | - | - |
| | Mother activity (cm/sec, log-transformed) | 0.126 | 0.072 | 3.085 | 0.090 | 1 | 0.000 |
| | Mother feeding (min) | 0.008 | 0.002 | 12.633 | 0.001 | 1 | 0.102 |
| | Grandmother ID [m2] | 0.148 | 0.118 | 0.783 | 0.467 | 2 | 0.006 |
| | Grandmother ID [m3] | -0.001 | 0.106 | | | | |
| | Offspring size at birth (mm) | 0.068 | 0.090 | 0.584 | 0.451 | 1 | 0.002 |
| | Mother size at parturition (cm) | -0.038 | 0.234 | 0.027 | 0.872 | 1 | 0.000 |
| | Observations | 34 | | | | | |
| | $R^2$ / $R^2$ adjusted | 0.391 / 0.256 | | | | | |
| | AIC | 39.596 | | | | | |
| Offspring feeding (min, brood average) | (Intercept) | 10.867 | 48.508 | - | - | - | - |
| | Mother feeding (min) | -0.090 | 0.126 | 0.505 | 0.483 | 1 | 0.018 |
| | Mother activity (cm/sec, log-transformed) | 1.408 | 4.215 | 0.112 | 0.741 | 1 | 0.004 |
| | Grandmother ID [m2] | -5.367 | 6.931 | 3.667 | 0.039 | 2 | 0.214 |
| | Grandmother ID [m3] | -15.542 | 6.188 | | | | |
| | Offspring size (mm, brood average) | 12.473 | 5.246 | 5.652 | 0.025 | 1 | 0.173 |
| | Mother size at parturition (cm) | -22.786 | 13.725 | 2.756 | 0.108 | 1 | 0.093 |
| | Observations | 34 | | | | | |
| | $R^2$ / $R^2$ adjusted | 0.404 / 0.272 | | | | | |
| | AIC | 262.857 | | | | | |

### Supplementary Information 3: Additional figures

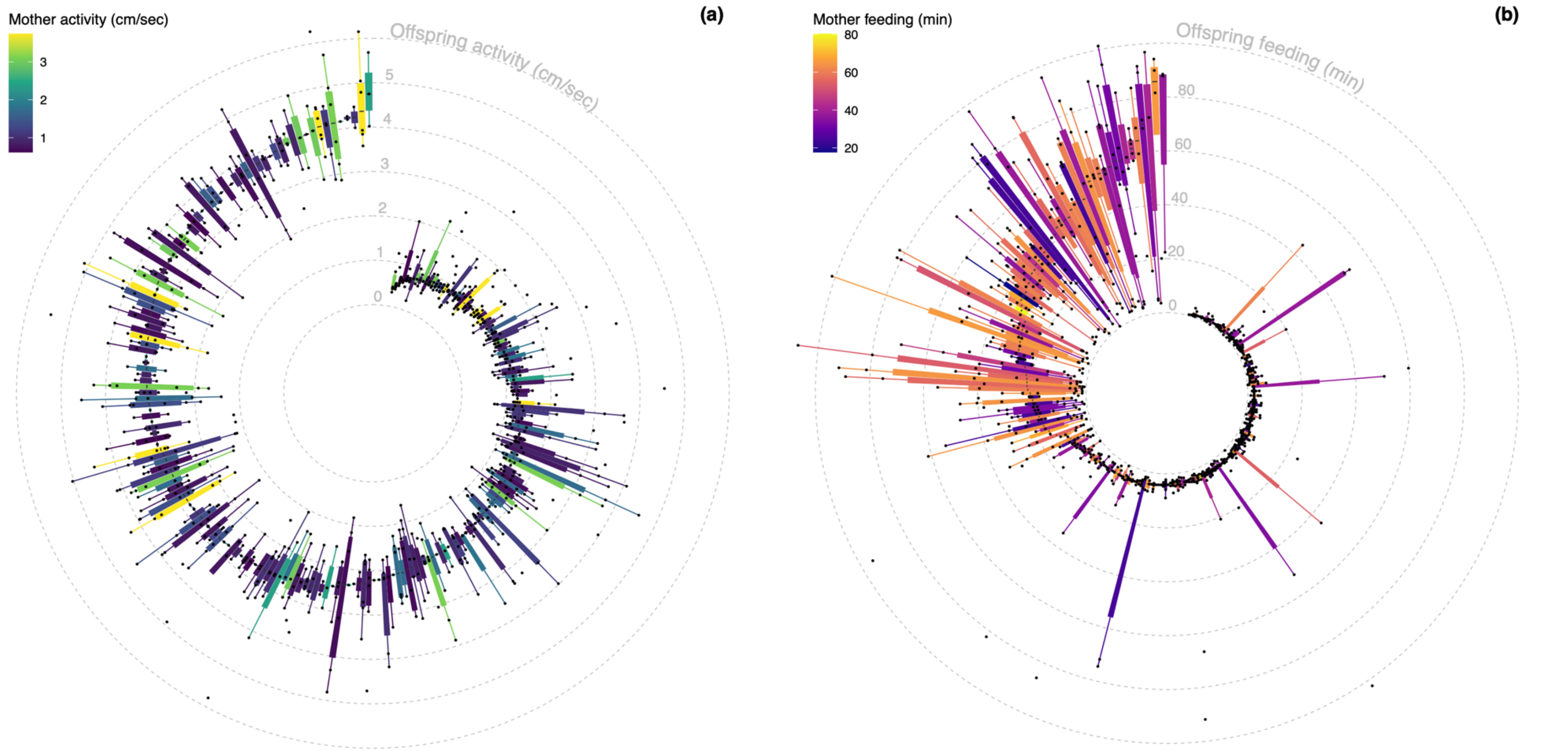

**Supplementary Figure S1**. Circular boxplot displaying daily (a) activity and (b) time spent feeding of *N* = 232 offspring from *N* = 34 mothers, observed over the first 2-7 days of life. (a-b) Shown are median (middle line), 25th to 75th percentile (box), and 5th to 95th percentile (whiskers) as well as the raw data (points) for each offspring (*N* = 922 daily behavioral measurements for offspring activity and feeding, respectively).

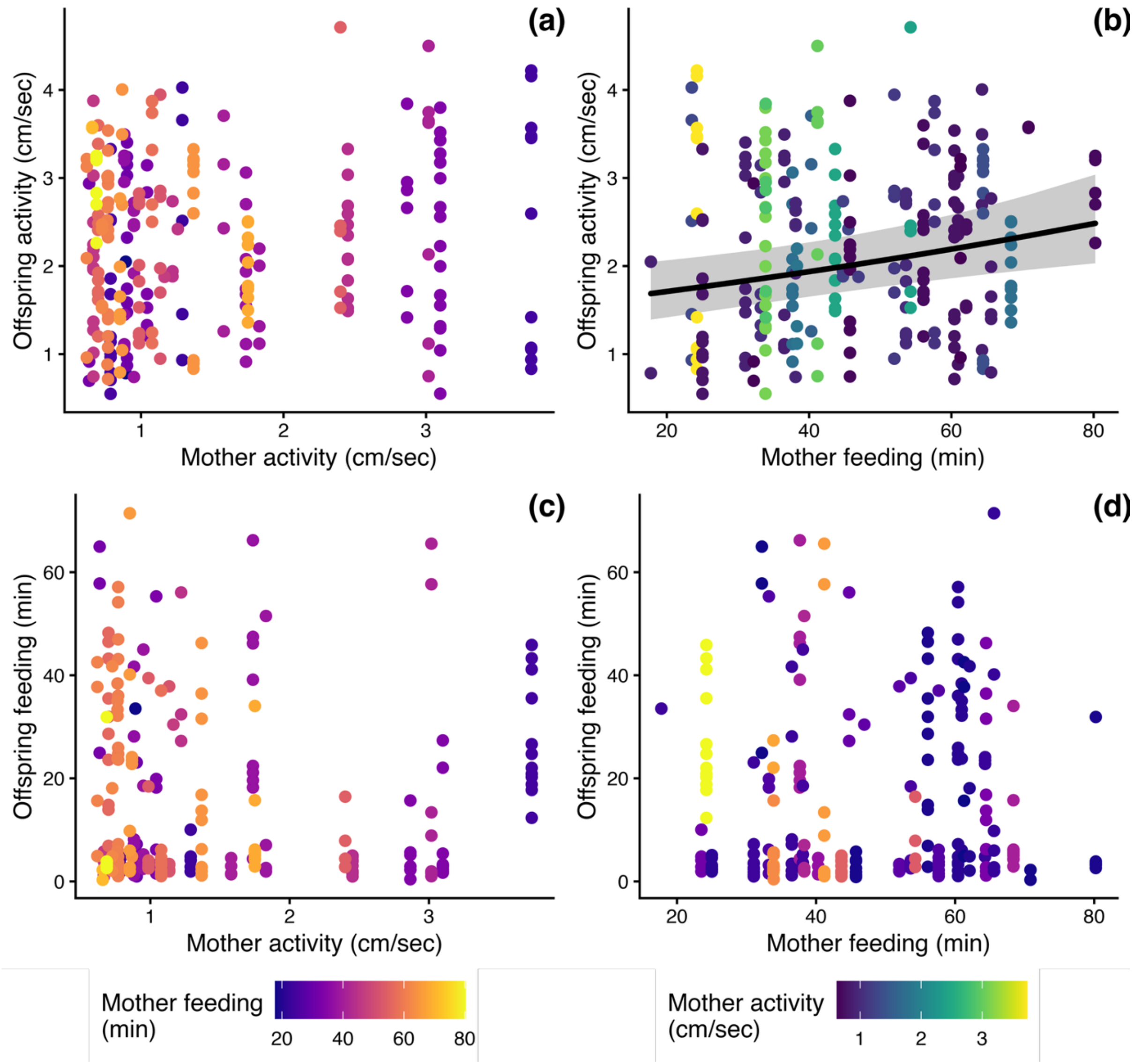

**Supplementary Figure S2**. Offspring activity shows a strong but statistically non-significant positive association with mother activity (a) and is positively associated with mother feeding behavior (b). In contrast, offspring feeding behavior is not associated with either mother activity (c) or mother feeding behavior (d). In contrast to Figure 1 in the main text, which presents aggregated mother-level means and standard deviations, this figure shows individual offspring means (N = 232 offspring) plotted against maternal means (N = 34 mothers). Activity values were log-transformed for statistical analyses and subsequently back-transformed for plotting to aid interpretability.

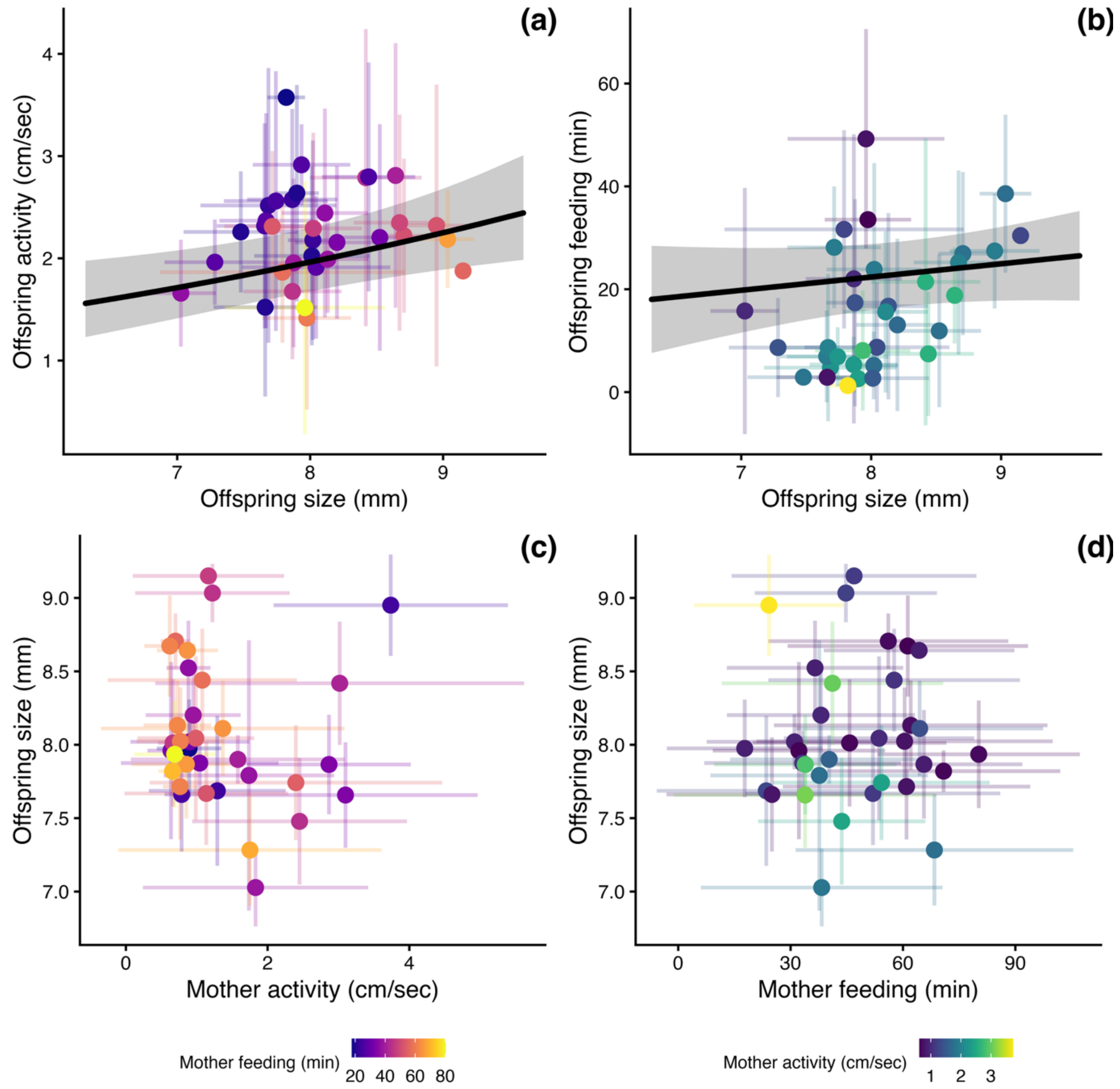

**Figure 3**. **Supplementary Figure 2.** Mother (a) activity and (b) feeding behavior do not predict offspring size. Offspring size, on the other hand, predicts both offspring (c) and (d) feeding behavior. (a-d) Data points represent the mean per mother (N = 34). Error bars reflect the standard deviation of daily maternal behavior (based on $N_{\text{maternal activity}}$ = 941 and $N_{\text{maternal feeding}}$ = 928 data points); standard deviation in mean offspring size at birth (based on *N* = 232 offspring), or the standard deviation in mean offspring behavior (based on *N* = 922 data points of daily offspring activity and feeding, respectively).

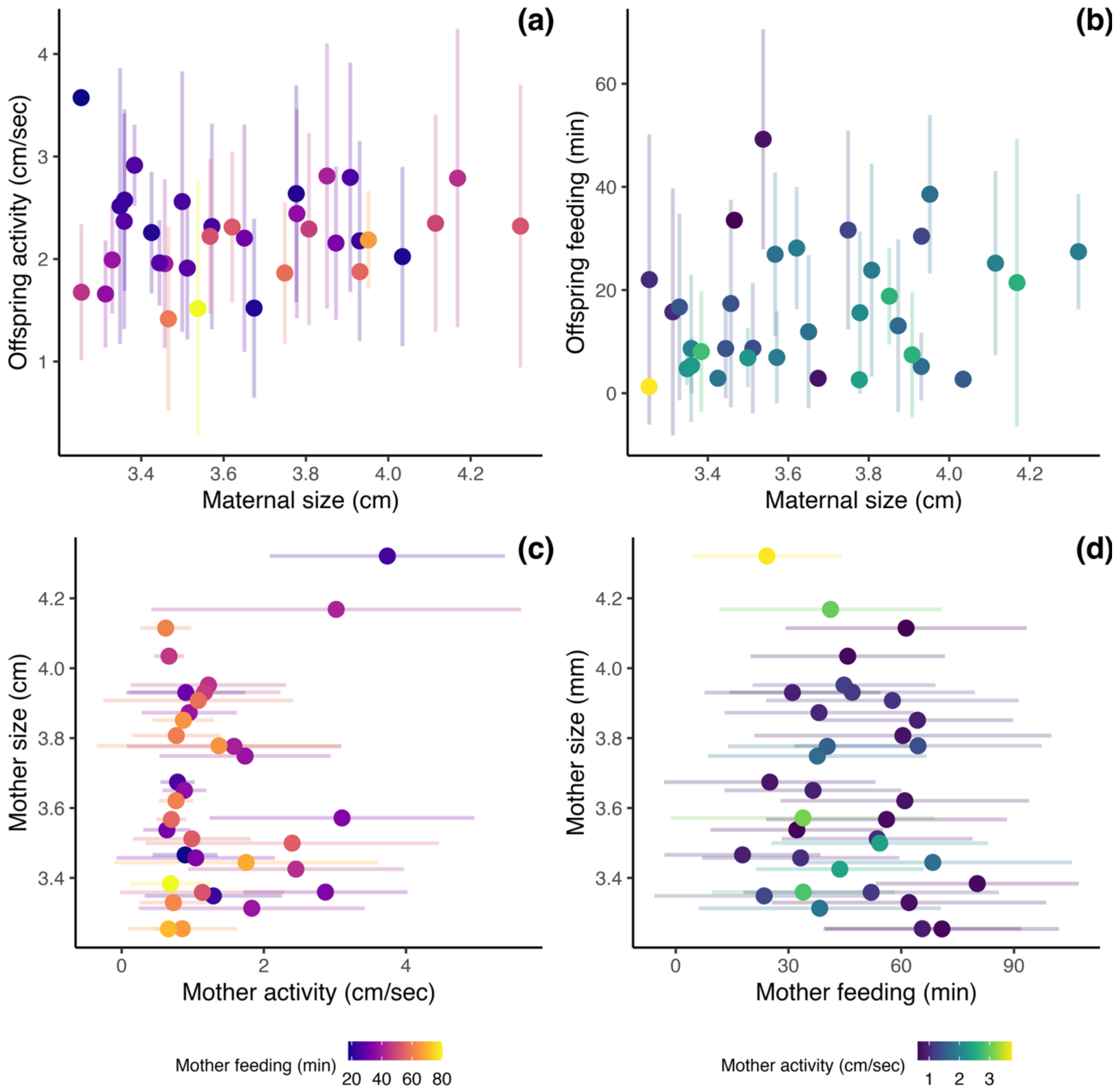

**Supplementary Figure 4.** Mother (a) activity and (b) feeding behavior do not predict maternal size at parturition. Offspring size, on the other hand, predicts both offspring (c) and (d) feeding behavior. (a-d) Data points represent the mean per mother (N = 34). Error bars reflect the standard deviation of daily maternal behavior (based on $N_{(\text{maternal activity})}$ = 941 and $N_{(\text{maternal feeding})}$ = 928 data points); standard deviation in mean offspring size at birth (based on N = 232 offspring), or the standard deviation in mean offspring behavior (based on N = 922 data points of daily offspring activity and feeding, respectively).

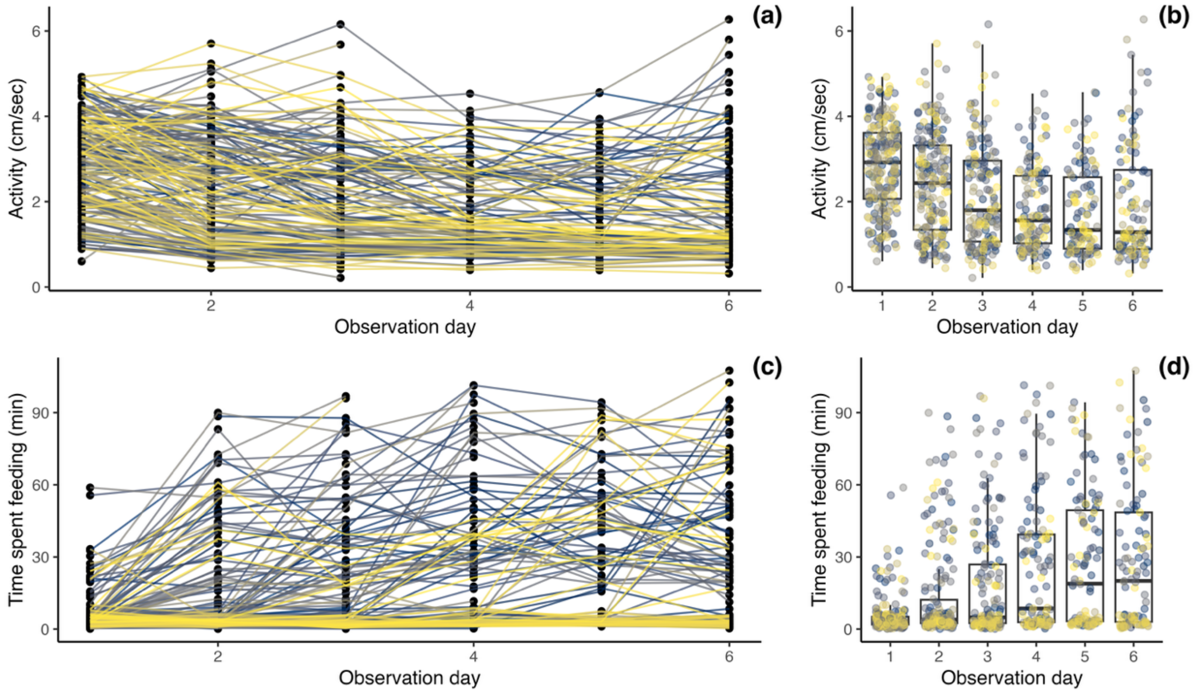

**Supplementary Figure 5**. (a-b) Offspring activity decreases over the first 6 days of life (linear mixed-effects model with daily activity as response, observation day as predictor, and offspring identity as random term, $N$ = 232 offspring from $N$ = 34 mothers: intercept ± SE = 2.911 ± 0.075, estimate ± SE = -0.216 ± 0.015, $\chi^2$ = 191.14, p-value < 0.001, $R^2$ = 0.097). (c-d) Offspring time spent feeding increases over the first 6 days of life (linear mixed-effects model with daily feeding time as response, observation day as predictor, and offspring identity as random term, $N$ = 232 offspring from $N$ = 34 mothers: intercept ± SE = 1.729 ± 1.480, estimate ± SE = -5.148 ± 0.345, $\chi^2$ = 195.69, p-value < 0.001, $R^2$ = 0.136). (a, c) Points represent individual, daily behavioral values, lines individual behavioral trajectories over observation days, coloration by ID. (b, d) Boxplots with median (middle line), $25^{th}$ to $75^{th}$ percentile (box), and $5^{th}$ to $95^{th}$ percentile (whiskers) as well as the raw data (points) for each offspring and day.

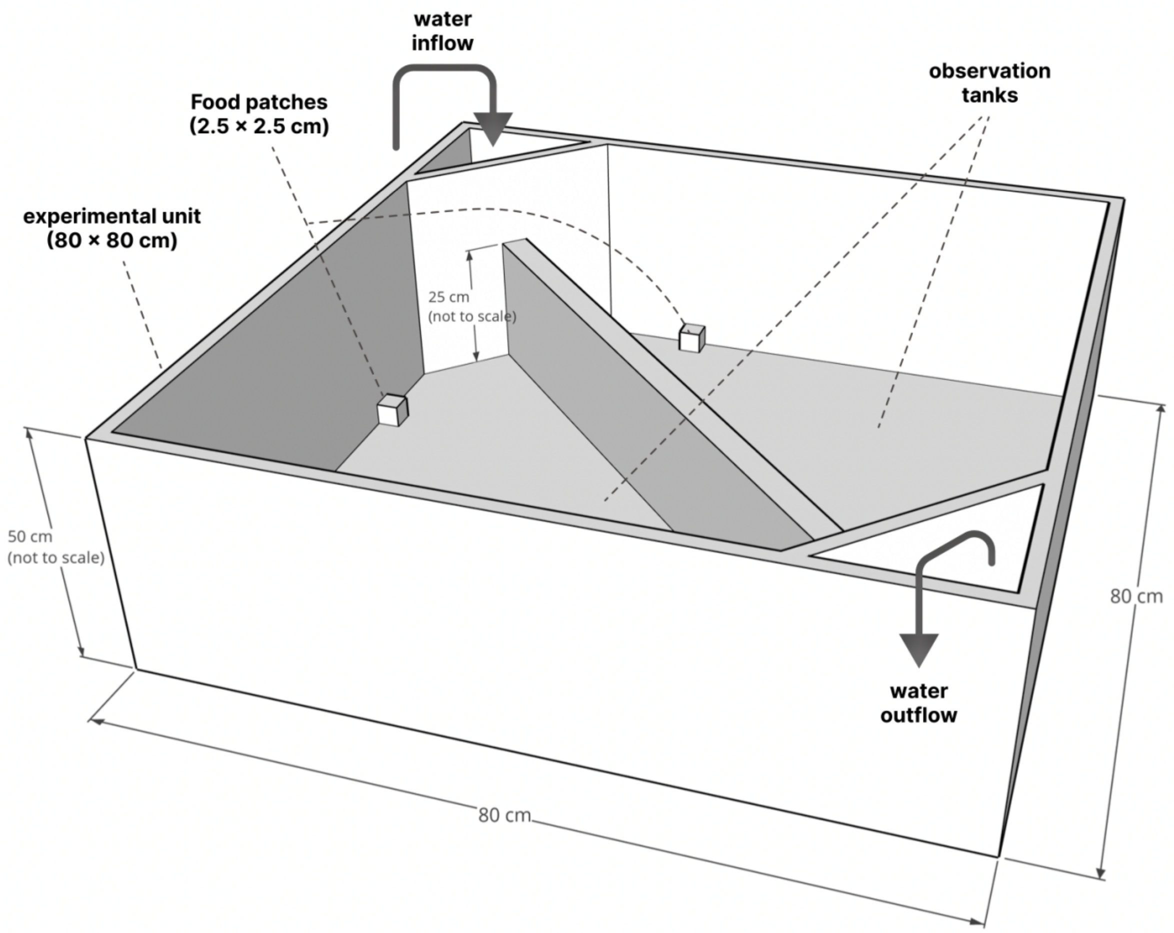

**Supplementary Figure 6**. Experimental set-up for behavioral observations. Figure and legend adopted from Scherer et al. (2023): "One experimental unit with 2 observation tanks. Water level in the tanks: 7 cm. Food patches were present during the feeding only. Observation tanks were illuminated individually from below with 4 LEDs per tank (each LED is 100cm, 12V, color temperature = 5500 K, light output = approx. 1570 lumen; tanks were manufactured from white polyethylene, which allowed light from underneath to get through). There was no visual contact between observation tanks, but tanks were connected via a flow-through water system (24 observation tanks split into 4 flow-through systems)."

### Supplementary Information 4: 9-Day behavioral assays

**Supplementary Table 17**. Results of linear models testing whether maternal behavior (activity, feeding) observed during the first 28 days of life predicts activity in a new tank (top), novel object response (middle), and sociability (bottom) in standard behavioral assays.

| **Response** | **Predictors** | **Estimates** | **SE** | **F** | **p** | **df** | **partial $R^2$** |
|---|---|---|---|---|---|---|---|
| Activity in a new tank (cm/sec, log-transformed) | (Intercept) | 0.203 | 0.208 | - | - | - | - |
| | Mother activity (cm/sec, log-transformed) | 0.581 | 0.130 | 19.889 | 0.001 | 1 | 0.407 |
| | Mother feeding (min) | 0.001 | 0.004 | 0.077 | 0.783 | 1 | 0.003 |
| | Grandmother ID [m2] | -0.427 | 0.205 | 2.869 | 0.073 | 2 | 0.165 |
| | Grandmother ID [m3] | 0.033 | 0.129 | | | | |
| | Observations | 34 | | | | | |
| | $R^2$ / $R^2$ adjusted | 0.465 / 0.391 | | | | | |
| | AIC | 49.807 | | | | | |
| Average distance to a novel object (cm) | (Intercept) | 28.612 | 2.572 | - | - | - | - |
| | Mother activity (cm/sec, log-transformed) | -1.035 | 1.613 | 0.412 | 0.526 | 1 | 0.014 |
| | Mother feeding (min) | -0.035 | 0.050 | 0.482 | 0.493 | 1 | 0.016 |
| | Grandmother ID [m2] | 2.206 | 2.538 | 2.494 | 0.100 | 2 | 0.147 |
| | Grandmother ID [m3] | -2.469 | 1.599 | | | | |
| | Observations | 34 | | | | | |
| | $R^2$ / $R^2$ adjusted | 0.184 / 0.072 | | | | | |
| | AIC | 199.598 | | | | | |
| Sociability (arcsine square root transformed) | (Intercept) | 1.366 | 0.072 | - | - | - | - |
| | Mother activity (cm/sec, log-transformed) | -0.069 | 0.045 | 2.357 | 0.136 | 1 | 0.075 |
| | Mother feeding (min) | -0.001 | 0.001 | 0.129 | 0.722 | 1 | 0.004 |
| | Grandmother ID [m2] | 0.019 | 0.071 | 0.591 | 0.561 | 2 | 0.039 |
| | Grandmother ID [m3] | 0.048 | 0.044 | | | | |
| | Observations | 34 | | | | | |
| | $R^2$ / $R^2$ adjusted | 0.098 / -0.026 | | | | | |
| | AIC | -43.967 | | | | | |

**Supplementary Table 18**. Results of linear models testing whether maternal behavior observed during standard behavioral assays (activity in a new tank, novel object response, and sociability) predict offspring activity (top) or feeding behavior (bottom).

| Response | Predictors | Estimates | SE | F | p | df | partial $R^2$ |
|---|---|---|---|---|---|---|---|
| Offspring activity (cm/sec, log-transformed; brood average) | (Intercept) | 1.147 | 0.507 | - | - | - | - |
| | Activity in a new tank (cm/sec, log-transformed) | 0.073 | 0.090 | 0.647 | 0.428 | 1 | 0.023 |
| | Average distance to a novel object (cm) | -0.010 | 0.009 | 1.302 | 0.264 | 1 | 0.045 |
| | Sociability (arcsine square root transformed) | -0.100 | 0.333 | 0.090 | 0.766 | 1 | 0.003 |
| | Grandmother ID [m2] | 0.191 | 0.117 | 1.423 | 0.258 | 2 | 0.092 |
| | Grandmother ID [m3] | -0.003 | 0.082 | | | | |
| | Observations | 34 | | | | | |
| | $R^2$ / $R^2$ adjusted | 0.141 / -0.012 | | | | | |
| | AIC | 49.291 | | | | | |
| Offspring feeding (min, brood average) | (Intercept) | -10.123 | 26.471 | - | - | - | - |
| | Activity in a new tank (cm/sec, log-transformed) | -2.778 | 4.707 | 0.348 | 0.561 | 1 | 0.012 |
| | Average distance to a novel object (cm) | 0.218 | 0.474 | 0.211 | 0.651 | 1 | 0.008 |
| | Sociability (arcsine square root transformed) | 20.889 | 17.393 | 0.090 | 0.241 | 1 | 0.049 |
| | Grandmother ID [m2] | -7.986 | 6.125 | 4.383 | 0.022 | 2 | 0.238 |
| | Grandmother ID [m3] | -12.401 | 4.285 | | | | |
| | Observations | 34 | | | | | |
| | $R^2$ / $R^2$ adjusted | 0.332 / 0.213 | | | | | |
| | AIC | 264.731 | | | | | |
